# Structure and evolutionary history of a large family of NLR proteins in the zebrafish

**DOI:** 10.1101/022061

**Authors:** Kerstin Howe, Philipp H. Schiffer, Julia Zielinski, Thomas Wiehe, Gavin K. Laird, John Marioni, Onuralp Soylemez, Fyodor Kondrashov, Maria Leptin

**Author notes:** ORCiD: 0000-0003-2237-513X. ORCiD: 0000-0001-6776-0934. ORCiD: 0000-0003-2294-0135. ORCiD: 0000-0001-9092-0852. ORCiD, 0000-0001-8308-6855. ORCiD: 0000-0001-7097-348X. All online supplementary material can be found on Figshare under the accession number http://dx.doi.org/10.6084/m9.figshare.1473092.

## Abstract

NACHT- and Leucine-Rich-Repeat-containing domain (NLR) proteins act as cytoplasmic sensors for pathogen- and danger-associated molecular patterns and are found throughout the plant and animal kingdoms. In addition to having a small set of conserved NLRs, the genomes in some animal lineages contain massive expansions of this gene family. One of these arose in fishes, after the creation of a gene fusion that combined the core NLR domains with another domain used for immune recognition, the PRY/SPRY or B30.2 domain. We have analysed the expanded NLR gene family in zebrafish, which contains 368 genes, and studied its evolutionary history. The encoded proteins share a defining overall structure, but individual domains show different evolutionary trajectories. Our results suggest gene conversion homogenizes NACHT and B30.2 domain sequences among different gene subfamilies, however, the functional implications of its action remains unclear. The majority of the genes are located on the long arm of chromosome 4, interspersed with several other large multi-gene families, including a new family encoding proteins with multiple tandem arrays of Zinc fingers. This suggests that chromosome 4 may be a hotspot for rapid evolutionary change in zebrafish.

## INTRODUCTION

The need to adapt to new environments is a strong driving force for diversification during evolution. In particular, pathogens, with their immense diversity and their ability to subvert host defense mechanisms, force organisms to develop ways to recognize them and keep them in check. The diversity and adaptability of pathogen recognition systems rely on a range of genetic mechanisms, from somatic recombination, hypermutation and exon shuffling, to gene conversion and gene duplication to generate the necessary spectrum of molecules.

The family of NACHT-domain (Koonin and Aravind 2000) and Leucine Rich Repeat containing (NLR) proteins reviewed in (Proell et al. 2008; Ting et al. 2008) act as sensors for sterile and pathogen-associated stress signals in all multicellular organisms. In vertebrates, a set of seven conserved NLR proteins are shared across a wide range of species. These are the sensor for apoptotic signals, APAF1, the transcriptional regulator CIITA, the inflammasome and nodosome proteins NOD1, NOD2, NOD3/NlrC3, Nod9/Nlrx1 and the as yet uncharacterized NachtP1 or NWD1 (Stein et al. 2007; Kufer and Sansonetti 2011). Other NLR proteins are shared by only a few species, or are unique to a species. Some non-vertebrates, such as sea urchins or corals, have very large families of NLR-encoding genes (Bonardi et al. 2012), but an extreme example of species-specific expansion can be found in zebrafish (Stein et al. 2007). Such species-specific gene family expansions suggest adaptive genome evolution in response to specific environments, most probably different pathogens (Liu et al. 2013).

The zebrafish has become a widely used model system for the study of disease and immunity (Rowe et al. 2014; Goody et al. 2014), and a good understanding of its immune repertoire is necessary for the interpretation of experimental results, for example in genetic screens or in drug screens. In a previous study we discovered more than 200 NLR-protein encoding genes (Stein et al. 2007). The initial description and subsequent analyses (Laing et al. 2008; van der Aa et al. 2009) have led to the following conclusions: The zebrafish specific NLRs have a well-conserved NACHT domain, with a ~70 amino-acid extension upstream of the NACHT domain, the Fisna-domain. This domain characterises this class of NLR proteins and is found in all sequenced teleost fish genomes. The NLR proteins can be divided into four groups, each defined by sequence similarity in the NACHT and Fisna domain that is greater than the sequence similarity shared across the whole family, and these groups also differ in their N-terminal motifs. Groups 1 and 2 have death-fold domains, groups 2 to 4 contain repeats of a peptide motif that is only found in this type of NLR protein. In the initial description, all of the novel NLR proteins ended with the Leucine-Rich-Repeats, but it was later found (van der Aa et al. 2009) that several of them had an additional domain at the C-terminus, an SPRY/B30.2 domain (PF00622). This domain also occurs in another multi-gene family implicated in innate immunity, the fintrims (van der Aa et al. 2009).

The initial identification of the genes and subsequent analyses by others (Laing et al. 2008) suffered from the limitations of the then available Zv6 assembly and gene annotations (published in 2006). This was not only caused by a limited amount of available data for long range assembly arrangements and supporting evidence for gene modeling - including the lack of well annotated homologues from other species - but also by the very high homology between the NLR genes and their clustered arrangement. Consequently, many assembly regions were collapsed and many of the gene models were incomplete. We have now revisited this gene family to improve the genome assembly in the regions of interest, to manually re-annotate and refine the gene structures and to provide a full description and analysis.

Our re-annotation and analysis of the NLR gene family in zebrafish shows a structured set of more than 400 genes with different evolutionary pressures on the different parts of the proteins, suggesting functional diversification between the groups.

## RESULTS

### 1. Identification of all NLR genes in the zebrafish genome

To identify the entire set of fish-specific NLR encoding genes in the zebrafish genome, we collected several lists of candidate genes based on the Zv9 assembly (GCA_000002035.2). We identified genomic regions containing domain motifs via hmmsearch (hmmer.janelia.org/search/hmmsearch), electronic PCR (Schuler 1997), TBLASTN searches, and by mining the existing annotations for keywords (Supplementary Methods). This collection was purged of gene models belonging to other known families, e.g. fintrims. We identified all overlapping Ensembl and VEGA gene models for the remaining regions of interest. The VEGA gene models were refined and extended through manual annotation and both gene sets merged, resulting in 421 NLR gene models.

Of these, 16 were not members of the family we had previously called ‘novel fish NLR proteins’ (Stein et al. 2007); these included the seven previously described NLR genes that are highly conserved among all vertebrates, and 9 other NLR genes that had a different structure from those described previously and below (Table 1). Of the remaining predictions for NLR proteins, which we will from now on refer to as NLR-B30.2 proteins, as explained below, 37 were pseudogenes, leaving 368 predicted protein-coding NLR genes (online supplementary material, Figshare: http://dx.doi.org/10.6084/m9.figshare.1473092). The genome assembly components carrying these gene models were checked for correct placement and relocated if necessary, thereby contributing to the improvements for the GRCz10 assembly (GCA_000002035.3).

**Table 1.**
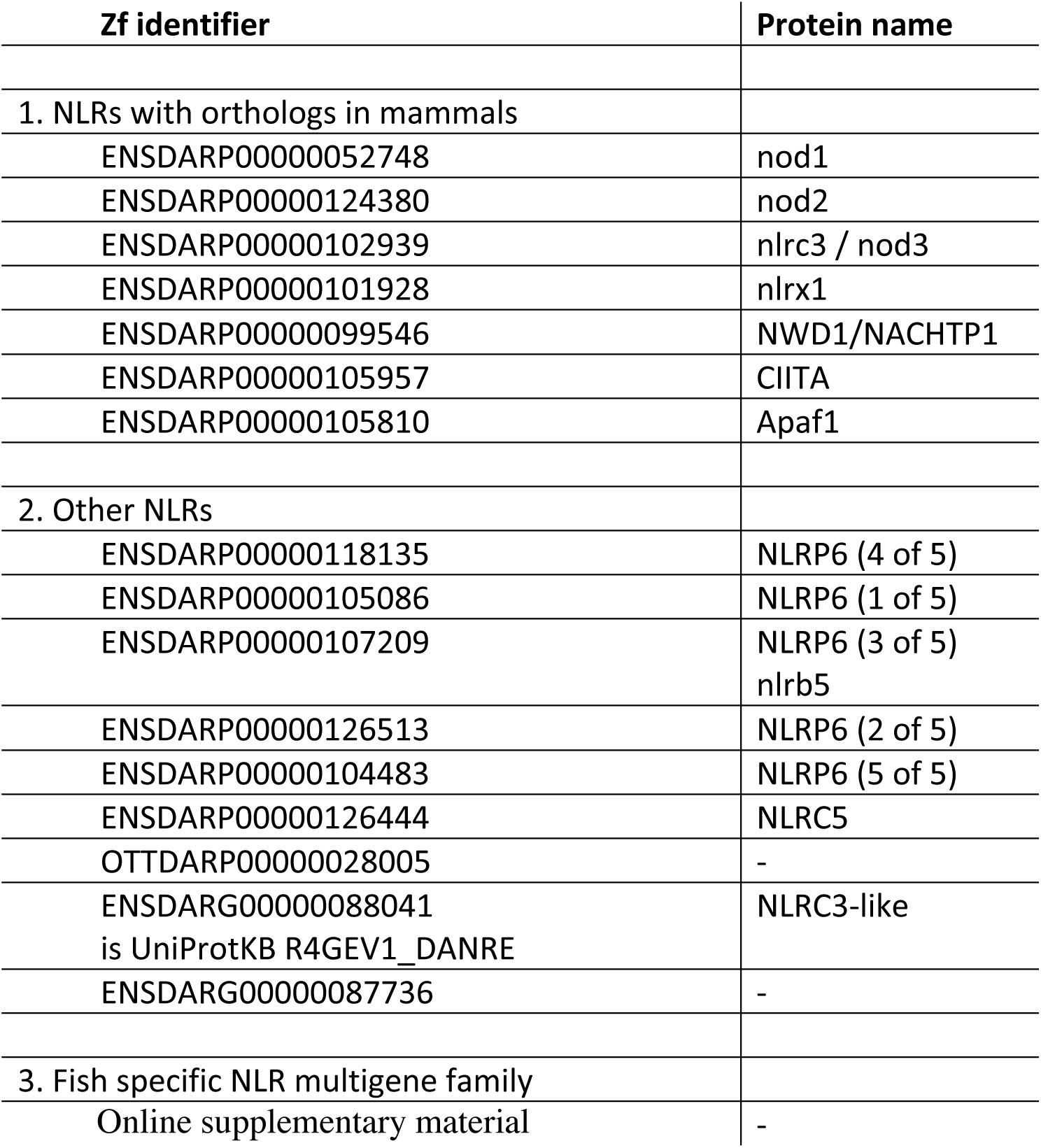
All NLR proteins in the zebrafish genome.

### 2. Structure, conservation and divergence

#### 2a. Domain structure of the NLR family members

The original set of 205 genes described in (Stein et al. 2007) had been divided into four groups based on sequence similarity in the Fisna and NACHT domains, and the sequence elements upstream of the Fisna domain. The current dataset shows that all four groups share the Fisna domain, the NACHT domain and the LRRs. A death-fold domain, a B30.2 domain and N-terminal peptide repeats are present in different subfamilies. Fig. 1 and Supplementary Fig. 1 show an overview of an alignment of the sequences and a schematic representation of the structure of the proteins, as discussed below.

**Figure 1:**
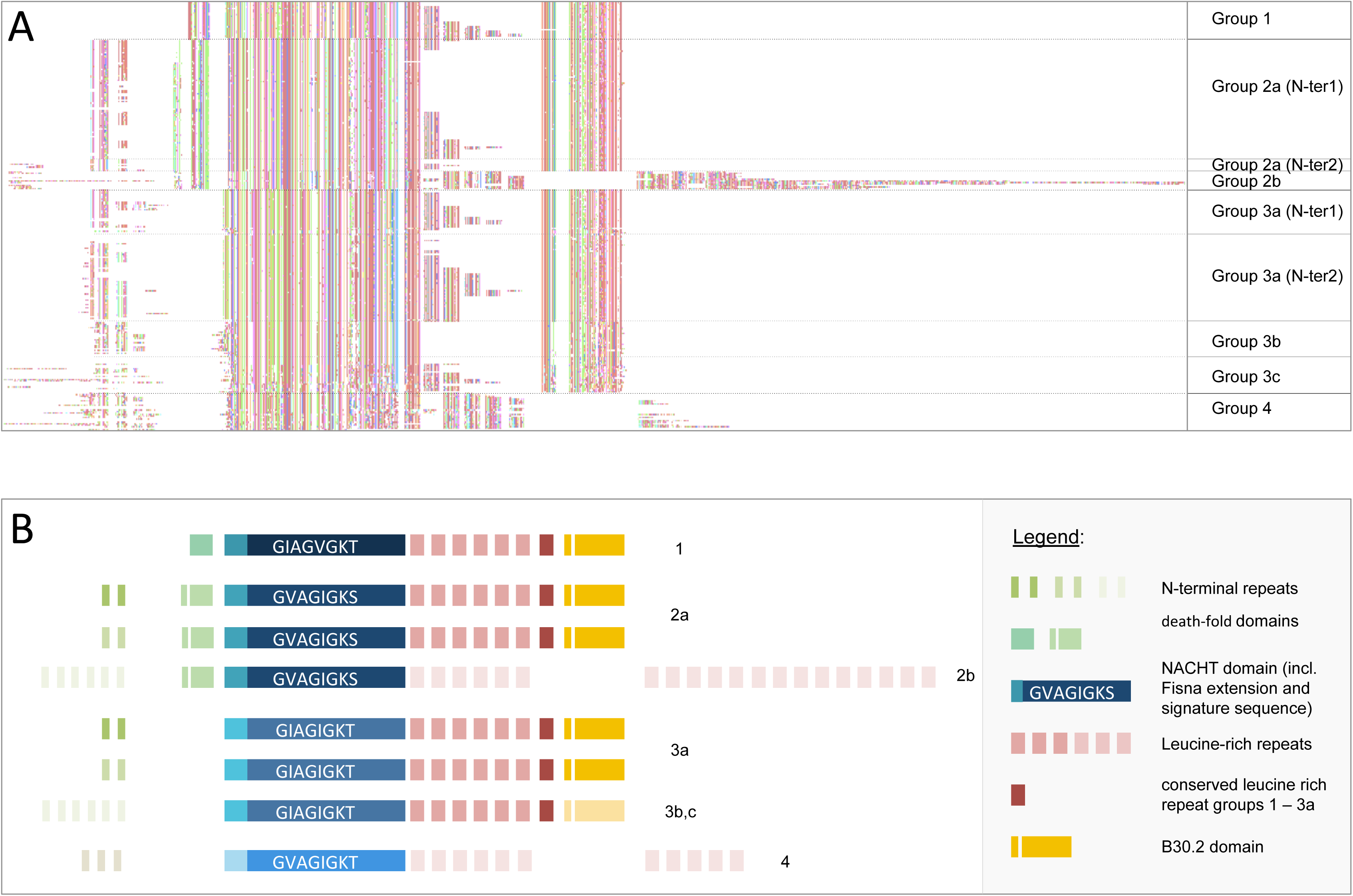
Structure of the fish-specific NLR-B30.2 protein family members. A. Alignment overview of 288 proteins for which full length predictions are available. Regions with long insertions in a few of the genes that had resulted in the introduction of long gaps in the alignment were deleted for the sake of clarity and simplicity. Gaps were manually introduced to highlight intron/exon boundaries, except between the C-terminal extensively duplicated LRRs in group 2b and the extensively duplicated N-terminal repeats in groups 2b and 3b. The colour coding is a random assignment of colours to amino acid created in Jalview. A gap was inserted between the LRRs 6 and 7 of groups 2b and 3b to allow the conserved C-terminal LRR and the B30.2 domains of groups 1, 2a and 3a to be positioned immediately after the 1-6 LRRs in these groups. B. An alignment of the full set of predictions is provided in the online supplementary material. A schematic representation of the protein domains in each group, on the same scale as in the alignment above. Each box represents an exon. Please refer to Supplementary Figure 2 and the alignment file (online supplementary material) for the many details not captured in this simplified figure.

##### Fisna and NACHT

The sequence similarities in the Fisna and the NACHT domain in our updated gene set confirm our previous subdivision into four main groups. A defining motif for each group is the sequence of the Walker A motif, for which the consensus over the whole set is G[IV]AG[IV]AGK[TS] and each group has its own, conserved motif (Fig. 1, Supplementary Figure 2). Some of the groups can be subdivided further. For example, group 2 consists of two subgroups (Figure 1, Supplementary Figure 1), one large and very homogeneous (group 2a), the other smaller, and also homogeneous, with all genes located in a cluster on chromosome 22 (group 2b). Group 3 has several subgroups that differ in their N-terminal peptides, or their LRRs, or their B30.2 domains (Fig 1.).

We previously found good matches for the Fisna-domain only in fish NLR proteins. Thus, the presence of this domain in addition to the particular Walker A motifs is a diagnostic property of the protein family. However, short stretches of this domain resembled peptides within mammalian Nod1 and Nod2 (Stein et al. 2007). The evolutionary origin of this part of the protein is unclear, but the fact that it is encoded in the same exon as the NACHT domain makes it unlikely to have originated through insertion of an exon from other genes.

We used the now larger collection of Fisna sequences to build an HMM for a Pfam family (PF14484) and to search for homologies within mammalian proteins. This revealed alignments with high significance to mammalian members of the NLR family (NLRP3 and NLRP12), with the matching sequence located immediately upstream of the NACHT domain. Secondary structure predictions based on two representatives from zebrafish and the rat using the PSIPRED workbench suggest that the zebrafish Fisna domain may take on the same conformation as the corresponding mammalian proteins (Fig. 2). Together, these findings suggest that contrary to previous assumptions, the Fisna domain was present in the common ancestor of mammalian and fish NLR proteins. Whether it forms a separate domain, or is simply an N-terminal extension of the NACHT-domain, similar to the additional helices seen in the structure of another NLR protein, Apaf1, remains to be determined by experimental work.

**Figure 2:**
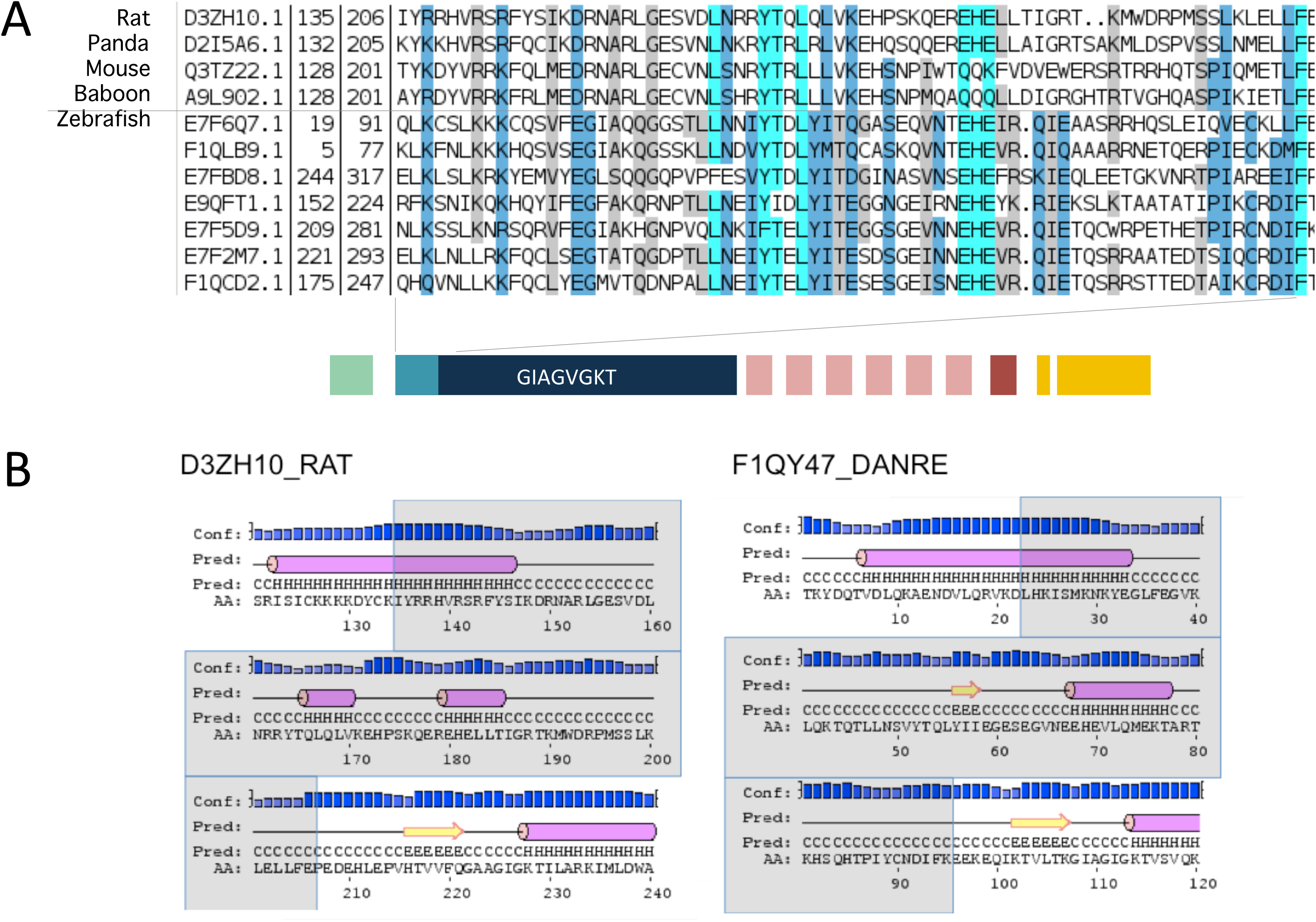
The Fisna domain and its homologs in mammalian proteins. A. Alignment of sequences identified in mammalian genomes using an HMM search with PF14484 and examples of zebrafish Fisna domains. B. Secondary structure predictions for selected examples.

##### Death-fold domains and N-terminal repeats

The four group-specific similarities continue upstream of the Fisna-domain: Groups 1 and 2 both have a death-fold domain; a Pyrin-domain (PYD) with an N-terminal peptide characteristic of BIR domains (http://elm.eu.org/elmPages/LIG_BIR_II_1.html) in the former, and a PYD-like domain in the latter. The predicted proteins of groups 2,3 and 4 contain several repeats of a ~30 amino acid peptide motif. The repeats are not all identical, but occur in two main types. Surprisingly, both of the main types of N-terminal repeat can associate with either group 2 or group 3 NACHT domains, indicating extensive exon-shuffling between family members.

##### LRRs

The proteins of all groups contain several Leucine Rich Repeats (LRRs; Pfam Clan: CL0022). The LRRs in groups 1, 2a and 3 are very similar to each other and occur in a similar pattern; each protein contains between 2 and 7 repeats, with each repeat consisting of two LRRs (Supplementary Fig. 3). There are two types of LRR repeat: the last LRR, immediately upstream of the B30.2 domain, occurs in each gene exactly once, and barely differs between the genes. The other type of LRR in groups 1, 2a and 3 can vary in number between 0 and 6, and these are more divergent in sequence. Thus, similar to the situation with the LRRs in the lamprey VLR proteins (Rast and Buckley 2013), the C-terminal LRR seems to be fixed, whereas the others vary more and are duplicated to varying degrees. Groups 2b and 4 do not show this arrangement, but they have yet another type of LRR, which can occur up to more than 20 times.

**Figure 3:**
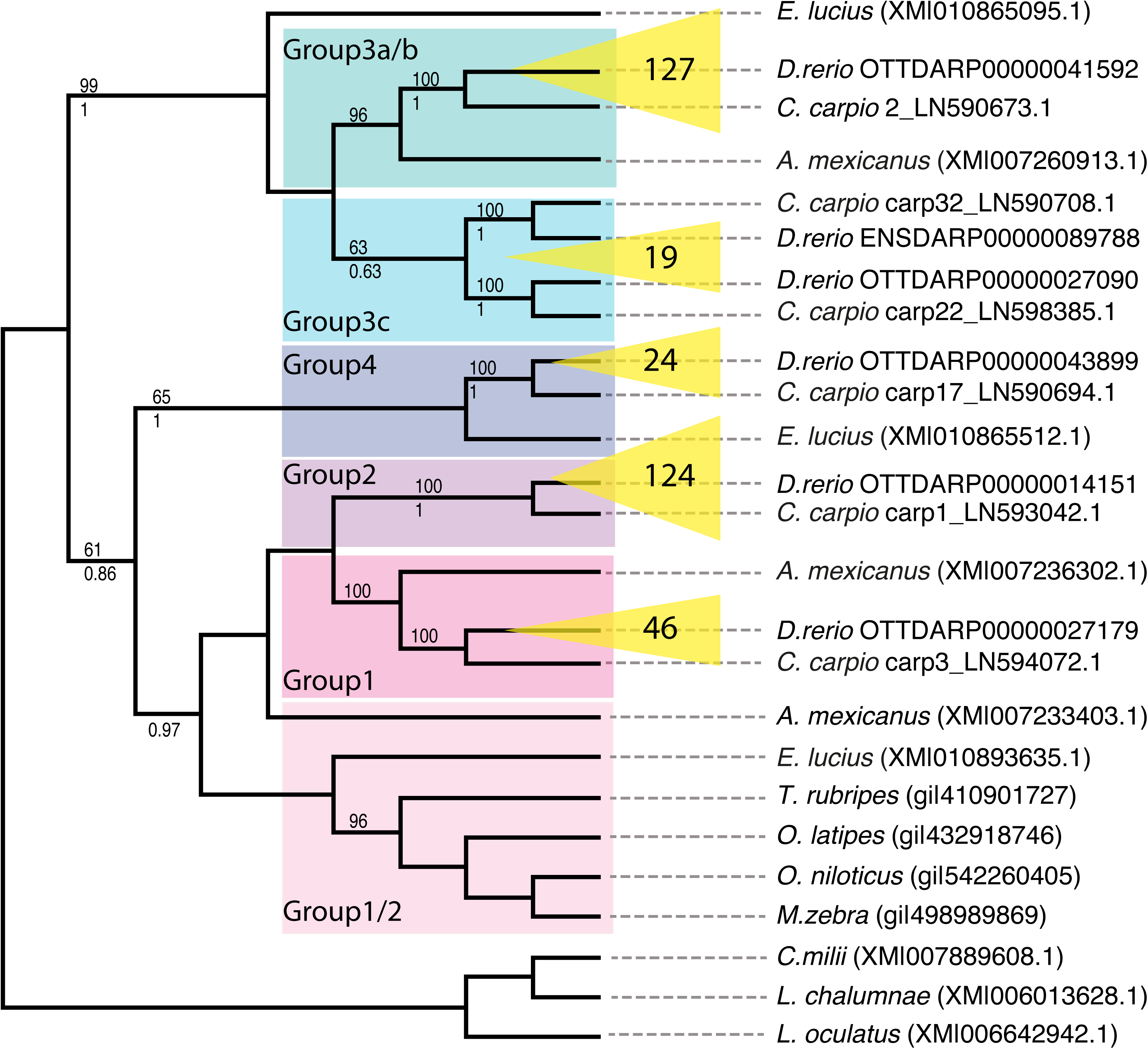
Relationships between NLR-B30.2 genes in different fishes. Phylogenetic tree resulting from a recursive phylogenetic analysis. Inflations of groups in zebrafish are indicated in yellow with numbers of genes per group displayed. Bootstrap values are given above each branch where higher than 50% and pp values (from MrBayes) below each branch, where they are higher than 50% and the topology is congruent.

##### SPRY/B30.2 domain

A B30.2 domain has so far been reported to be present in some but not all fish NLR proteins (van der Aa et al. 2009). Our current set of sequences shows that the presence of the B30.2 domain is restricted to groups 1, 2a and 3, and the domain is missing in groups 2b and 4 (Fig. 1, supplementary Fig. 5). In view of the extreme similarity between the N-terminal parts (NACHT and death-fold domain) of groups 2a and 2b, and the overall conservation of the gene structure throughout the whole family, it seems most likely the that B30.2 domain was present in the common ancestor of the family, but lost by groups 2b and 4, rather than independently gained by the other groups. We therefore refer to the entire family as the NLR-B30.2 protein family.

#### 2b. Intron-exon structure

All genes in this family have the same exon-intron structure (Supplementary Fig 2). Briefly, each domain is encoded on a single exon. The largest exon contains the NACHT domain, including the winged helical and superhelical domains, as well as the N-terminal Fisna domain.

#### 2c. Sequence variation between the groups

Visual inspection of the aligned amino acid sequences of all four groups showed that some parts of the proteins were highly conserved, while others showed many amino acid substitutions. The NACHT domains of groups 1, 2a and 3a show little sequence divergence within each group, but strong divergence between the groups (Supplementary Figure 3). In contrast, there is no recognizable group-specific sequence pattern in the B30.2 domain. Instead, a number of apparently sporadic substitutions are seen throughout the entire set (Fig. 1). Also the last LRR is almost completely invariant between groups 1, 2a and 3a and the other LRRs do not show group-specific variations. Thus, on the one hand, proteins with different NACHT domains share similar B30.2 domains, and on the other, proteins with nearly identical NACHT domains and N-terminal motifs, such as those in groups 2a and 2b, have different C-termini. Furthermore, there are two major types of N-terminal repeats, but rather than each being associated with a specific group, each can be combined with either group 2a or group 3a (Supplementary Fig. 1: group 2a (N-ter1) and group 2a (N-ter2)). Both the shuffling of N-termini and the unequal divergence of the NACHT and B30.2 domains suggest a complex evolutionary history of the gene family.

### 3. Evolutionary history

#### 3a. Evidence for gene conversion within the four zebrafish NLR-B30.2 groups

The high similarity of the amino acid sequences of the NACHT domains within groups and the clear distinction between groups suggest a possible functional need for this separation of families and perhaps a directed evolutionary process that creates this situation.

To test this, we calculated rates of synonymous sequence divergence (dS). We studied only those groups that show high intra-group conservation of the NACHT domains (groups 1, 2a, 3a and 3b). We considered groups 3a and 3b separately because inspection of the protein alignment suggested that although they were classified as one group by virtue of their NACHT domain, their B30.2 domains appeared more diverged. We left out-group 3c, as its NACHT domains are more divergent, and may represent further groupings. This left 220 zebrafish NLR-B30.2 genes with full-length sequences covering both domains. We considered synonymous divergence (dS) separately for the NACHT and the B30.2 domains within and between each of the four clusters. For each of the 220 genes we obtained the pairwise synonymous divergences (dS) to all other 219 genes and within each group recorded the lowest dS to sequences from all 4 groups (Table 2).

**Table 2.** dS calculations for NACHT and B30.2 domains within and between groups and
subgroups.

| NACHT | 1 | 2a | 3a | 3b |
| --- | --- | --- | --- | --- |
| 1 | **$0.05 \pm 0.006$** | $1.14 \pm 0.011$ | $1.11 \pm 0.010$ | $0.96 \pm 0.011$ |
| 2a | $1.27 \pm 0.001$ | **$0.01 \pm 0.000$** | $1.11 \pm 0.001$ | $1.26 \pm 0.001$ |
| 3a | $1.16 \pm 0.001$ | $1.11 \pm 0.001$ | **$0.01 \pm 0.000$** | **$0.09 \pm 0.000$** |
| 3b | $1.02 \pm 0.004$ | $1.31 \pm 0.003$ | **$0.09 \pm 0.001$** | **$0.02 \pm 0.002$** |
| B30.2 |  |  |  |  |
| 1 | **$0.035 \pm 0.002$** | **$0.036 \pm 0.004$** | **$0.039 \pm 0.004$** | $0.466 \pm 0.006$ |
| 2a | **$0.039 \pm 0.000$** | **$0.019 \pm 0.000$** | **$0.025 \pm 0.000$** | $0.467 \pm 0.000$ |
| 3a | **$0.042 \pm 0.001$** | **$0.028 \pm 0.001$** | **$0.026 \pm 0.001$** | $0.471 \pm 0.000$ |
| 3b | $0.671 \pm 0.006$ | $0.551 \pm 0.005$ | $0.600 \pm 0.005$ | $0.217 \pm 0.006$ |

The patterns of synonymous divergence between groups were clearly different for NACHT and B30.2 domains and confirmed the different behaviours of the two domains seen we had noted from the alignment of the protein sequences.

The rate of synonymous sequence substitutions in the NACHT domains was very low when comparing members within any one group (0.01 - 0.05); we find values comparable to the divergence observed among the closest of any sister species in the vertebrate lineage. The values for comparisons between groups were 20- to 100-fold higher. The only case where the between-group comparison also gave a low value was for group 3a versus 3b, confirming their classification by NACHT domain as one group.

The B30.2 domains showed a different pattern. The dS values ranged between 0.02 and 0.04 for groups 1, 2a and 3a, and the values for the between-group comparisons were minimally or not at all higher than for within-group comparisons. Therefore, the B30.2 domains of these three groups have not significantly diverged from each other. The B30.2 domain sequences of the members of group 3c were divergent from each other as well as diverged from B30.2 domains from genes in groups 1, 2a and 3a (Table 2).

This different pattern of synonymous divergence of the two domains between groups is most parsimoniously explained by on-going gene conversion in the NACHT-B30.2 gene family. This confirms the conclusion derived from inspection of the amino acid sequences, and indicates that the NACHT domains have been homogenized within the individual groups, whereas the B30.2 domains have been homogenized between groups 1, 2a and 3a.

#### 3b. Origin of the NLR-B30.2 gene groups

The degree of apparent gene conversion within the NLR-B30.2 gene family makes it difficult or impossible to judge when the groups arose or when the amplification of each group occurred during fish evolution. To find out whether the groups arose in the zebrafish or in an ancestral species, we compared the NLR-B30.2 genes in the zebrafish with those in the closest relative for which a whole genome sequence exists, the carp, as well as NLR-B30.2 genes in other vertebrate genomes (see Materials and Methods).

To investigate when the subdivision into groups first occurred, we performed a recursive phylogenetic analysis of the NLR-B30.2 genes from available high quality fish genomes: the northern pike *Esox lucius*, the spotted gar *Lepisosteus oculatus*, the Mexican tetra *Astyanax mexicanus*, the pufferfish *Takifugu rubripes*, the cichlid *Maylandia zebra*, the Japanese rice fish *Oryzias latipes*, the Nile tilapia *O. niloticus* and the common carp *Cyprinus carpio*, as well as the coelacanth *Latimeria chalumnae*, with the elephant shark *Callorhinchus milii* as outgroups.

This analysis showed that the split into groups occurred before the zebrafish-carp divergence (Fig. 3). Groups 1, 2, 3a/b and 4 have clear orthologous relationships between carp and zebrafish: For example, group 2 from zebrafish is most closely related to a group of genes in the carp that is clearly distinct both from other carp NLR-B30.2 genes and from the other zebrafish genes.

Not unexpectedly, group 3c, which is more complex as judged by comparison of the sequences in the zebrafish, also shows a more complex evolutionary history. It falls into two groups, each of which has an orthologous group in the carp.

*A. mexicanus* and *E. lucius* each have groups of genes that cluster with groups of the zebrafish genes, rather than with each other, but not every group is represented in each of the species. Nevertheless, this suggests that the split is even older than the carp-zebrafish split, having perhaps occurred in basal teleosts. Sequences from *L. oculatus, L. chalumnae* and *C. milii* do not fall into these groups, showing that the split into groups must have occurred in the Clupeocephala.

In summary, the groups did not diverge independently from duplicated ancestral genes in each species, but in a common precursor. By contrast, the majority of genes within each group in each species arose by species-specific, independent amplification of a founding family member.

#### 3c. Co-occurrence of the NACHT and B30.2 domains

As first reported by (van der Aa et al. 2009), the NLR-B30.2 domain fusion is found in all teleost fish. Our collection of NLR-B30.2 genes showed that this domain fusion is older than the common ancestor of teleosts, as it was also present in *L. oculatus* (for example ENSLOCG00000000593), for which the genome sequence was not previously available. The NLR-B30.2 proteins predicted in *L. oculatus* also contain an N-terminal Fisna extension, though only distantly related to those in the teleosts, indicating that the ancestral gene included this extension. This is consistent with the fact that the N-terminal extension of the mammalian NLRP3 proteins has recognizable similarity to the Fisna domain, as described above.

We do not find evidence of the NLR-B30.2 fusion in any of the tetrapod genomes, nor in the *L. chalumnae* or *C. milii* genomes. These results indicate that the fusion occurred at least in the common ancestor of the Neopterygii, prior to the third whole genome duplication in the teleost lineage. Genome sequence data from sturgeon, paddlefish or bichir clades, which are currently not available, would provide further information on the point of emergence of the NLR-B30.2 fusion.

#### 3d. Expansion of the NLR-B30.2 family in the fish

The phylogeny of the NLR-B30.2 gene family also provides information on the timing of the lineage expansion of the NLR-B30.2 genes observed in the zebrafish. The C. carpio genome contains a similarly large family of NLR-B30.2 proteins, but due to the polyploidy of the species some of the nearly identical sequences may rather constitute alternative alleles rather than paralogues. *A. mexicanus*, a direct outgroup to the zebrafish-carp clade, features the second largest NLR-B30.2 gene family with ~100 members, showing that the lineage expansion began prior to the zebrafish-carp split. Other fish species, including the spotted gar have fewer than 10 NLR-B30.2 genes, while *E. lucius*, a Euteleost, has ~50. Thus, the initial gene expansion either occurred in the basal branches of Teleostei with a subsequent loss in some lineages, or independently in several lineages. Independent expansions and losses are a likely scenario, given the expansions of NLR genes in many other species, such as sponges and sea urchins (Yuen et al. 2014; Lapraz et al. 2006). The results on fish show that the expansion of the NLR-B30.2 family began as soon as the NLR-B30.2 fusion occurred, with different dynamics in different lineages.

#### 3e. Age of the NLR-B30.2 family relative to conserved NLRs

We compared the origin of the NLR-B30.2 gene family to the evolutionary history of other NLR genes. There are many species-specific expansions of NLRs, such as the Nalp proteins in mouse and human and the NLR-B30.2 genes in fish, as well as independent inflations in Amphioxus, sea urchin, and sponge. However, there are also the seven NLR proteins that are conserved in all vertebrates and show orthologous relationships. We collected the available orthologues of the conserved NLRs from key metazoan species and created an alignment. We also included all NLR genes from fish that did not belong to the NLR-B30.2 group (listed above and in M&M).

We found that two of the conserved vertebrate NLR genes appear to be shared by all animals (Fig. 4, Supplementary Fig. 5, and online supplementary material). The genes for NWD1 (first described in zebrafish as NACHT-P1) and Apaf1, must have been present in the last common ancestor of bilaterians and non-bilaterians, as they are found in sponges, cnidarians, and all bilaterians analysed. We could not find any candidates in comb jellies (ctenophores). The other five conserved NLR proteins - Nod1, Nod2, NLR3C, CIITA and NLRX1 – arose later in evolution, at the base of the gnathostomes. An additional gene, NLR3c-like, was present at this point, but appears to have been lost in the tetrapod lineage.

In summary, all of the conserved vertebrate NLRs are older than the NLR-B30.2 family. They are never duplicated and certainly not expanded to higher gene numbers in any species.

**Figure 4:**
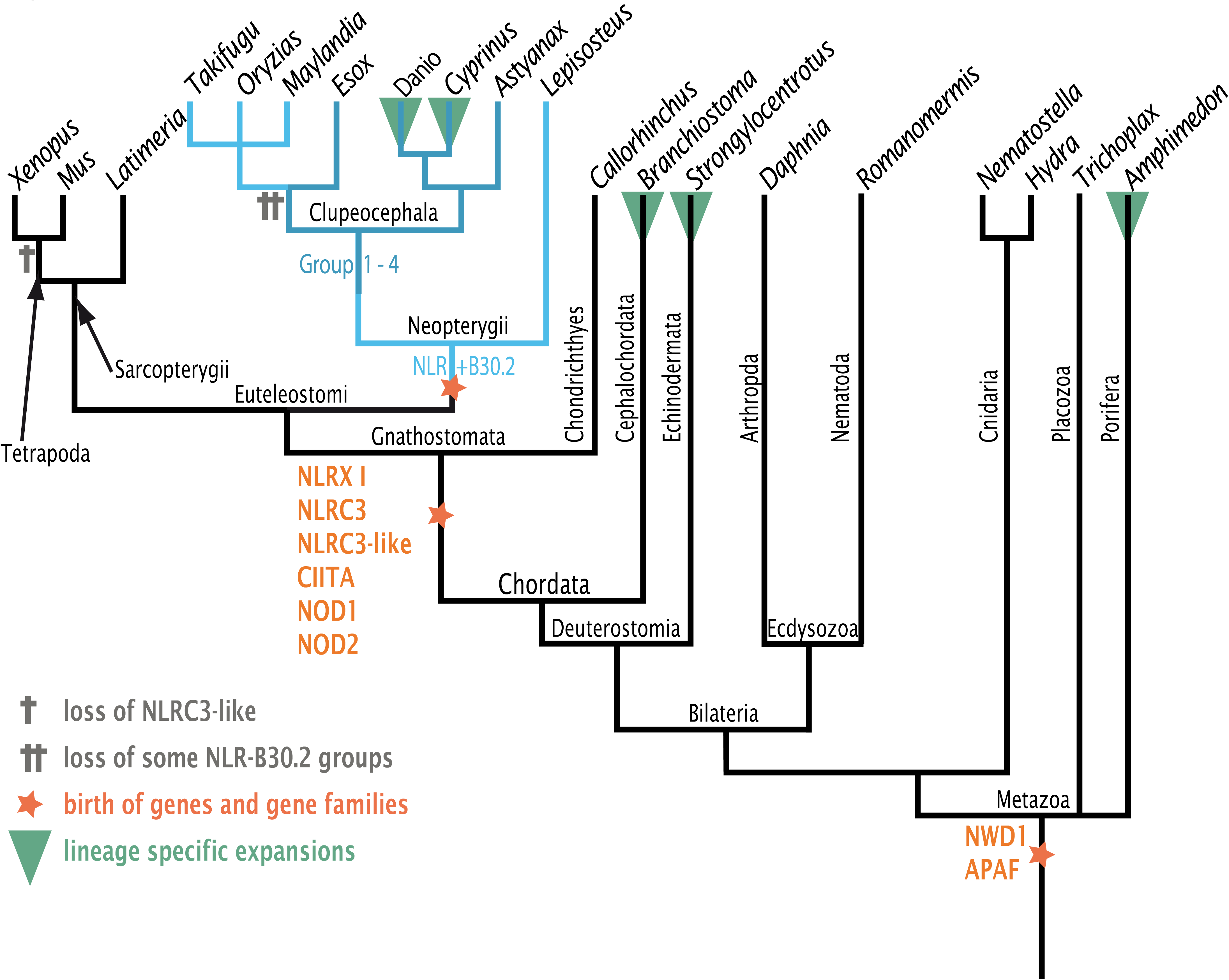
Evolutionary history of NLR genes. A reduced dendrogram of Metazoa based on the NCBI taxonomy database displaying key events in the evolution of NLR genes as described in the main text. See Supplementary Figure 6 and the supplementary online material for phylograms.

Important events in the evolutionary history of the NLR-B30.2 family and conserved NLR proteins are summarized in Figure 4 and Supplementary Figure 5. The oldest metazoan NLR proteins are NWD1 and Apaf1, while Nod1, Nod2, NLR3C, CIITA and NLRX1 and NLR3C-like were acquired in the lineage leading to gnathostomes. A fusion of the NLR and B30.2 domains then occurred in the Neopterygii lineage and subsequently these genes diversified into groups, within which they continued to expand in various fish lineages. The high similarity of the NACHT domains within the groups appears to have been maintained by gene conversion. Large-scale amplifications of NLR genes occurred independently in many species.

### 4. Genomic location

During the first survey of NLR genes on the zebrafish genome assembly Zv6, the genes were located on 22 different chromosomes, with some enrichment on chromosome 4 (50 genes) and chromosome 14 (47 genes) (Stein et al. 2007). Since this analysis, the assembly of the zebrafish genome has been significantly improved and the current Zv9 NLR gene set shows a more restricted distribution (Fig. 5), with 159 (44%) of the genes located on the right arm of chromosome 4. The remaining genes are distributed between 12 other chromosomes (153 genes) and unplaced scaffolds (56 genes).

**Figure 5:**
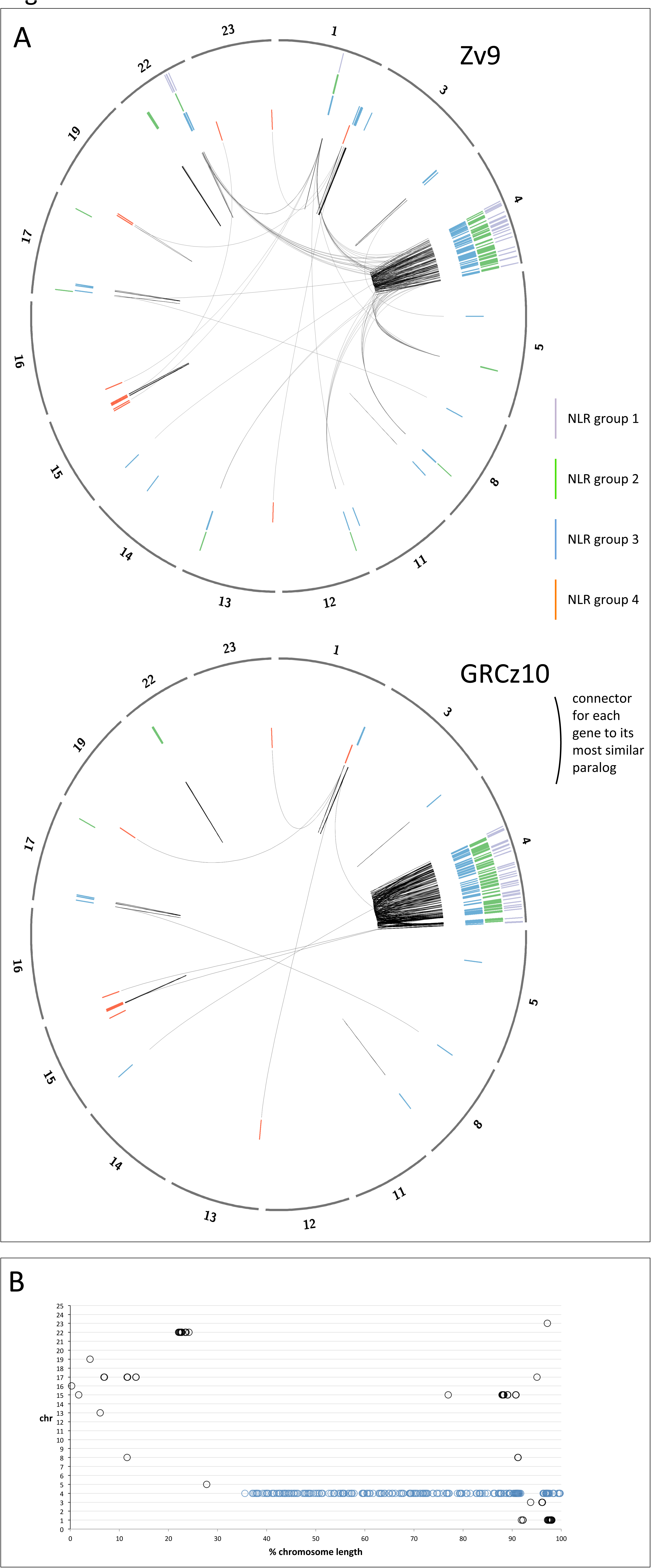
Location of the NLR genes in the genome in assemblies Zv9 and GRCz10. A. The chromosomes containing NLR-B30.2 genes are shown in the outer circle (note that corrections of the genome between Zv9 and the GRCz10 have changed the lengths of some of the chromosomes. The genes were annotated on Zv9 and lifted over to the GRCz10 path where possible as the GRCz10 gene set did not become available until May 2015. The members of the four groups of NLR genes are shown as radial lines within the circles with group 1 in the outermost and group 4 in the inner ring. Each gene is connected by a black line to its most closely related paralog, based on the number of amino acid substitutions per site calculated in MEGA5 (Poisson correction model). Most genes are most closely related to a near neighbour. The changes in the assembly have have lead to many genes that were closely related but resided on different chromosomes in Zv9 being located in closer proximity in GRCz10. B. Normalised location of NLR-B30.2 genes on chromosomes. Each chromosome is shown as a horizontal line of 100% length, and the NLR genes are plotted at their relative positions along the chromosome. Apart from the genes on chromosome 4 (marked in blue), all other genes are found within the first or last quarter of the chromosome.

Additional sequencing and data gathering by the Genome Reference Consortium since the release of Zv9 led to the rearrangement of multiple assembly components, including relocation of sequence to different chromosomes. These placements are based on manual curation by the Genome Reference Consortium, supported by genetic mapping data, clone end sequence placements and optical mapping data (Howe and Wood 2015).

The latest assembly, GRCz10, reveals the majority of the genes clustered on the long arm of chromosome 4, where 240 (66%) of the NLR-B30.2 genes, including all group 1 and group 2a genes, now reside (Fig. 5), whilst group 2b genes are now found exclusively in a cluster on chromosome 22. This gives an indication of the evolutionary history of these groups, and argues for group 2b having arisen by local duplication from one precursor gene in this location that had lost its B30.2 domain. Similarly, group 3 has its main group 3a on chromosome 4, with groups 3b and 3c on chromosomes 1 and 17, respectively, again as local clusters. Group 4 genes are found mostly on chromosome 15, some on chromosome 1, and they are excluded from chromosome 4. Both group 2 and group 3 have a few individual genes dispersed over other chromosomes; careful inspection of the evidence on which this allocation is based revealed no indications that it is incorrect. Some of the group 3 members on other chromosomes are more divergent from the consensus for this group, suggesting they may indeed have separated from the group early.

Within chromosome 4, no clear pattern can be detected in the distribution of the genes. We are, however, aware of possible shortcomings in the assembly of the long arm of chromosome 4; the highly repetitive nature of the sequence makes it difficult to exclude with absolute certainty shuffling of gene locations. In addition to containing multiple copies of 5S ribosomal DNA (Sola and Gornung 2001), 53% of all snRNAs and the majority of the NLR-B30.2 genes, chromosome 4 also contains multiple copies of genes encoding a particular type of Zn-finger protein, which we discuss below.

Finally, another striking feature of the distribution of the genes across the genome is that they tend to accumulate near the ends of the chromosomes, if we ignore the special situation of chromosome 4 (Fig. 5). With the exception of the cluster on chromosome 22, and two single genes on Chromosomes 5 and 15, all other genes (82% of the NLR genes outside chromosome 4) are located within 15% of chromosome ends.

### 5. Distribution of Fintrim and multiple Zn-finger encoding genes

We noticed that the NLR-B30.2 genes on chromosome 4 were often interspersed with genes encoding multiple tandem Zn-finger proteins. In some cases, gene models had been made which joined B30.2 domains with Zn-fingers, but our analyses showed that the B30.2-encoding exons instead belonged to a neighbouring NLR gene, rather than the more distant Zn-finger encoding exons. A possible explanation for the mis-annotation is that the predictions created apparent Fintrim genes. Fintrim proteins are composed of multiple Zn-fingers combined with a B30.2 domain and are assumed to act as sensors for immune stimuli (van der Aa et al. 2012). They are often found in the vicinity of, or at least on the same chromosome as, genes that are located in the MHC in mammals (MHC-associated genes are found not in one cluster in the zebrafish, but on four different chromosomes [3, 8, 16, 19]). We therefore analysed the distribution of the NLR-B30.2 genes relative to the location of fintrims and multiple Zn-finger encoding genes.

To establish a list of fintrim genes from the current assembly, we collected and refined candidate genes from the Zv9 genes set, resulting in 61 TRIM, 40 ftr and 18 btr genes (online supplementary material). An alignment of 283 B30.2 domains from the NLR@B30.2 genes and 117 from the fintrim and btr families shows that most of the B30.2 domains from the NLR@B30.2 genes are more closely related to each other than to those of the TRIM families. We find that there is no close association in the genome between the fintrim and the NLR@B30.2 genes (Fig. 6B), except for two cases where a single fintrim gene is found near an NLR@B30.2 gene cluster (chr1 and chr15). If anything, fintrim genes are excluded from regions where NLR@B30.2 genes are found and vice versa. The NLR@B30.2 genes are also not located in the vicinity of the regions in which the MHC@derived genes map in the current assembly.

**Figure 6:**
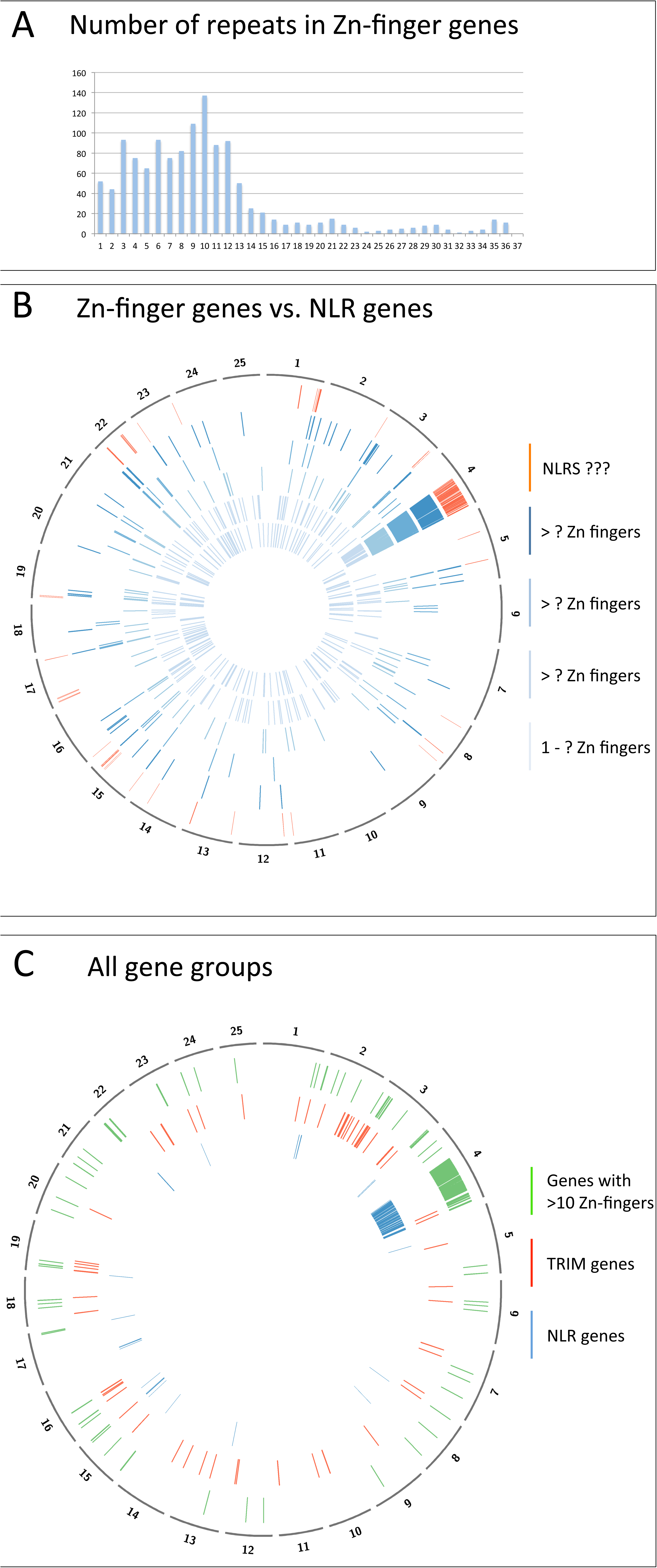
Genomic positions of genes for multi-Zn-finger proteins, Fintrims and NLR-B30.2 proteins. A. Frequency of genes encoding proteins with the indicated number of Zn-finger domains. Total number of Zn-finger encoding gene predictions: 1102. B. The genomic locations of Zn-finger encoding genes is plotted on a circos diagram representing the Zv9 assembly for five subgroups defined by the number of Zn-finger domains. Inner circles (light to dark blue): 1-3, 4-5, 6-7, 8-9, 10-12 and more than 12 domains; Outer circle (red): NLR genes C. Spatial distribution of NLR genes, TRIM genes and genes with at least 10 Zn-finger domains in the GRCz10 assembly (for gene liftover see Fig. 5).

However, as noted above, there were genes encoding multiple Zn-fingers interspersed among the NLR-B30.2 genes. Unlike the TRIMs, which contain C3HC4 (RING) and Znf-B-box domains (IPR001841 and IPR000315, respectively), the genes we had noticed on chromosome 4 encoded yet uncharacterised proteins consisting exclusively of tandem repeats of Zn-fingers of the classical C2H2 type (IPR007087). To investigate this new gene family further, we collected all gene models encoding Zn-fingers of this type. A total of 1259 gene models were found, with the number of repeats per gene ranging from 1 to 36. Some of the cases of genes with extremely large numbers of C2H2 domains (e.g. ENSDARP00000109314) may be mis-annotations that combine adjacent, shorter genes, which we did not analyse manually in further detail.

The encoded proteins with small numbers of Zn-fingers included many known proteins, including the Sna and Opa transcription factor families. These were broadly distributed in the genome and largely excluded from chromosome 4 (Fig. 6). By contrast, those with larger numbers of Zn-finger domains are progressively clustered in restricted regions of the genome. The majority are found on the right arm of chromosome 4, where they are interspersed in an irregular pattern among the NLR-B30.2 genes. Outside chromosome 4, some multi-Zn-finger genes co-locate with subsets of the TRIM genes, for example on chromosomes 3, 16 and 19, while others are located in regions where neither NLR-B30.2 genes nor TRIMs are found.

Thus the local duplications that may have led to the expansion of the NLR-B30.2 genes on chromosome 4 may also have duplicated the multi-Zn-finger genes, which have subsequently been transposed to other chromosomes.

## DISCUSSION

### Phylogeny of vertebrate NLR proteins

The family of NLR-B30.2 genes has been shaped by different genomic and genetic mechanisms at different times during evolution. These include repeated gene amplifications, shuffling of exons and gene fusions, gene conversion and positive selection for diversity. From the vertebrate viewpoint, the NLR genes can be divided into those that are present with orthologous relationships in all vertebrate genomes, and those that are shared by few or specific to single species.

The oldest NLR genes appear to be those encoding the ancestors of two conserved NLRs, Apaf1 and NWD1, which we find in all animal lineages. These proteins have not been reported to have immune functions. Apaf1, originally discovered as CED-4 in *C. elegans*, is an ancient regulator of apoptosis. The so far only function reported for NWD1, first identified in the zebrafish genome as NACHTP1 (Stein et al. 2007), is its involvement in androgen signalling in the context of prostrate cancer (Correa et al. 2014). It will be interesting to learn whether this is a special case of a more general immune function yet to be discovered, or whether, like Apaf1, this old gene does not have immune functions.

The other conserved genes first appear at the base of the gnathostomes, and all have roles in immunity or inflammation - whether as transcription factors or as inflammasome components.

In parallel, NLR genes have duplicated and often undergone extensive expansions independently in many species throughout evolution. This is the case, for example, for the members of the Nalp/NLRP family in the mouse and the NLR-B30.2 family we discuss here. The largest of the known early expansions were in the sponge *A. queenslandica*, the sea urchin *S. purpuratus* and the lancelet *B. floridae*, with about 120, 92, and 118 genes respectively (Yuen:2014hia; Huang et al. 2008), and it is likely that as more genomes are sequenced, more NLR expansions will be discovered. In vertebrates, the largest expansions are those of the NLR-B30.2 family, although we also find other NLR gene families, for example in the elephant shark *C. milii* (Supplementary Figure 5, online supplementary material).

The expansion of the families argues in favour of their involvement in immunity or broader stress reactions, as seen in numerous other examples. Expansions can increase the amount of gene product, for example to adapt to stressful environmental conditions (Kondrashov et al. 2002; Kondrashov 2012), as in the cold adaptation in several gene families expressed in Antarctic notothenioid fishes (Chen et al. 2008). Expansions can also increase the rate of sequence divergence between copies (Lynch 2002), thereby creating the variety of sequences that is needed especially for immune recognition, as in the case of antibodies and T-cell receptors, or the more recent example of the VLR genes in lampreys and hagfish (Li et al. 2013).

After their initial creation through the fusion of the NLR and B30.2 components, and initial duplication events, which the available data do not allow us to trace, the NLR-B30.2 genes diversified into groups. While these groups are recognizable in most of the fishes we analysed, not all fishes have all four groups. We see almost no divergence at all in the zebrafish genes between the NACHT domains within each group. The remarkably low rate of synonymous substitutions within the individual groups makes it likely that they are being kept distinct by constant gene conversion.

Gene conversion is not uncommon in gene families involved in immunity (see Pasquier, 2006, for review). It can create diversity, for example in antibodies e.g. (Wysocki and Gefter 1989) or in the MHC (reviewed in Martinsohn et al., 1999), but it can also homogenize genes, e.g. in the T cell receptor family (Jouvin-Marche et al. 1986). In the NLR-B30.2 family both outcomes are seen: within-group homogenisation in the NACHT domain, and creation of variation in the B30.2 domain.

The following evolutionary scenario for the zebrafish NLR-B30.2 gene family is the most parsimonious. Starting from few ancestral paralogous genes, conversion acted across the genes on both the NACHT and B30.2 domains. At some point, the NACHT domain appears to have become more prone to loss of gene conversion than the B30.2 domain.

Coupled with continuing gene duplications this may have led to the emergence of distinct groups largely defined by sequence divergence of the NACHT domain, while within each group gene conversion continued to homogenize the NACHT domain sequence. By contrast, the slower rate of loss of gene conversion in the B30.2 may have led to the maintenance of gene conversion in that domain across groups that have lost it for the NACHT domain.

It is worth speculating about the functional and selective forces that prevent sequence homogenization between the NACHT domains of different groups. If the proteins form large multimeric complexes, as the known inflammasome NLRs do, then their efficient functioning might require that only proteins from the same group can multimerize, for instance to elicit distinct down-stream signaling events. This is supported by the groups featuring different N-terminal domains. A mixed multimer may not be able to assemble a functional N-terminal effector complex.

The C-terminal domains – LRRs and B30.2 - do not show the same clear subdivision into families as the N-terminal and the NACHT domains, and homogenizing gene conversion must therefore have affected only part of each gene, or affected different parts differently. This is not without precedent, since gene conversion often proceeds across DNA segments of limited length (see Innan 2009 for review) and parts of a gene can escape sequence homogenization (Innan 2009; Teshima 2004).

Both LRRs and B30.2 domains have been implicated in recognition of pathogen- or danger-associated molecular patterns. The B30.2 domain of Trim5a binds to HIV-1 and is involved in blocking HIV-1 proliferation in monkeys (Stremlau et al. 2004). In Nod1 and Nod2, the LRRs are implicated in recognising components of pathogens (Inohara et al. 2005), including flagellin monomers (Akira et al. 2006), while in toll like receptor proteins they recognise viral dsRNA (Kawai and Akira 2009).

The sequences of the LRRs in the NLR-B30.2 genes are not particularly variable, and it therefore seems unlikely that they have a role in specific ligand-recognition. The B30.2 domains however show significant amino acid variation between the members. The related B30.2 domain in the fintrim genes has been suggested to be under positive selection to allow variation in specificity for pathogen recognition, making it likely for the B30.2 domain in the NLR-B30.2 proteins to have the same function. The gene conversion between the B30.2 domains across groups 1a, 2a and 3a that is suggested by the dS analysis may be a mechanism by which the diversity is created. It is conceivable that the acquisition of the B30.2 domain and the option to use it for specific recognition of a wide range of pathogens drove the amplification of these genes.

In summary, NLR-B30.2 gene family can be regarded as a product of mutation-driven evolution, where gene conversion as neutral process acting on a previously diversified paralogues maintains diversity, which then became the substrate of selection. While we lack sufficient genomic data from salt water fishes, we are tempted to speculate that the massive inflation of the NLR-B30.2 group may have occurred along with the adaptation to fresh water environments where fish would have encountered a greater abundance of pathogens (see e.g. Johnson and Paull, 2010).

### Shuffling between genes and creation of new genes

A further mechanism in the creation of the NLR-B30.2 family appears to have been exon shuffling, both within the family and between the NLR genes and other gene families. For example, the N-terminal peptide repeats occur in several variants, but a given variant is not strictly associated with any particular group: at least two of the variants are found both in association with group 2a and group 3.

We also find evidence for recombination with other immune genes. The B30.2 domain of the NLR-B30.2 proteins resembles most closely that of the fintrim proteins, a fish-specific gene family for which the origin of the fusion between the Zn-fingers with the B30.2 domain is not known (van der Aa et al. 2009), suggesting that exon shuffling occurred during the generation of the ancestral genes of the NLR-B30.2 and the fintrim gene families.

Apart from this possible case of exon-exchange, the relationship between the three large and partially related families – the NLR-B30.2 genes, the fintrim genes and the multi-Zn-finger genes we describe here – are unclear. While it is striking that the fintrims share the B30.2 domain with the NLR-B30.2 genes and the Zn-fingers with the multi-Zn-finger proteins, they do not preferentially map to the same regions of the genome, and the Zn-finger is of a different type. By contrast, the multi-Zn-finger genes are mostly found on chromosome 4, interspersed between the NLR-B30.2 genes. We have not attempted to trace the relative evolutionary histories of these three families.

A further gene that may have arisen from domain shuffling between these gene families is the human gene encoding pyrin (marenostrin/MEFV). Pyrin is a protein that is composed of an N-terminal PYD domain, for which the best match in the zebrafish is the PYD domain in the group 1 NLR-proteins. The C-terminal part of pyrin contains a Zinc-finger and a B30.2 domain, which resembles the zebrafish fintrim proteins of the btr family. The most likely interpretation for the origin of this gene, which must have arisen at the base of the tetrapods, is therefore a recombination between an NLR gene and a neighbouring fintrim gene.

### Chromosome 4

The zebrafish chromosome 4 has unusual properties. Its long arm is entirely heterochromatic, replicates late and shows a reduced recombination rate. It contains an accumulation of 5S rRNA, snRNA, tRNA and mir-430 clusters (Anderson et al. 2012; Howe et al. 2013), as well as the expanded protein coding gene here.

Chromosome 4 was recently shown to function as the sex chromosome in wild zebrafish ZW/WW sex determination, with the sex determining signal being located immediately downstream of the NLR-B30.2 expansion region (Wilson et al. 2014). The sex determination region in the grass carp may also be associated with NACHT domain encoding genes (Wang et al. 2015). This was concluded from the comparison of the genome sequences of one male and one female carp, where those regions present in the male and absent in the female were interpreted as sex-determining. In addition to the NACHT domain genes, this region also included other immunity genes, such as the immunoglobulin V–set, ABC transporters, and proteasome subunits. While the co-location between sex determination and immune signalling molecules we describe here may support this conclusion, it is of course equally possible that the finding in the grass carp is simply caused by allelic diversity in these highly variable genes between the two individuals. It is nevertheless intriguing that two fast evolving genetic systems are located in such close proximity in zebrafish. Perhaps, after an initial round of NLR gene duplications, a run-away evolutionary process of further amplification set in to create the present chromosome 4, which is now a hotspot for rapid evolutionary processes.

## METHODS

### Re-annotation of NLR genes in the zebrafish genome

To establish a complete list of all genes encoding NLR proteins in the zebrafish genome, we first conducted a search of the Zv9 genome assembly for sequences that encoded the characteristic protein domains, using a combination of approaches. We constructed a hidden Markov model (HMM) for the Fisna domain and used this together with the HMM for the NACHT domain obtained from PFAM to search the Zv9 assembly with hmmsearch (hmmer.janelia.org/search/hmmsearch), resulting in 297 Fisna and 328 NACHT locations (see online supplementary material). As an alternative way to identify NACHT domains specific for the novel NLRs, we ran electronic PCRs (PMID: 9149949) with primer sets for a segment stretching from the C-domain into the winged helix domain that we had used for experimental analysis of the genes (unpublished work). Each set of primers was specific for one of the NLR groups (Supplementary Methods). This resulted in 321 hits. To find regions in the genome encoding B30.2 domains we conducted a TBLASTN search, which yielded 503 hits. As B30.2 domains also occur in other large, immune-related protein families (see below), such a high number of domains was consistent with expectations.

Secondly, we collected all Ensembl genes overlapping the above motifs (487 predicted genes) and also all manually annotated genes (vega.sanger.ac.uk) that had been marked as NLR or as containing a NACHT domain during manual annotations in the past (307 predicted genes).

The collection was purged of gene models that did not match the criteria for being novel NLRs, excluding e.g. the B30.2 domain-containing fintrim genes. Sixteen NACHT domain proteins in the combined list do not belong to the group of novel fish NLRs since they do not contain the Fisna-domain, and the sequence surrounding their WalkerA motifs does not match the one typical for the novel NLRs. They include the seven conserved NLRs that are orthologous across all vertebrate species (Nod1, Nod2, Nlrc3, Nlrx1, CIITA, Apaf1, NWD1/NachtP1), and nine further proteins with an NLR structure. (Table 1 and online supplementary material)

Comparison of the purged gene sets with the genomic regions that encoded parts of NLR proteins showed that many genes in this family had been annotated incorrectly, and for others there were no predictions at all, probably due to the repetitive nature of this gene family and the limited availability of supporting evidence in the form of cDNAs.

The regions containing the sequences identified in our searches were therefore re-annotated manually, correcting and adding gene models to create full-length genes. This re-annotation had to be restricted to regions located on finished sequence, since whole genome shotgun (WGS) contigs in Zv9 were not accessible to manual annotation. For these contigs, the automated Ensembl gene models were retained in their original form, recognizable in our final list by their “ENSDARG” identifier (Supplementary Excel Table1). The resulting protein sequences were then aligned using Clustal-Omega (Sievers et al. 2011) or Muscle (Edgar 2004) and compared. Sequences that appeared truncated were analysed further by searching for additional exons to complete them, until, in an iterative process, we had optimised them. Some sequences remained incomplete, either because they were located next to sequence gaps, or because no additional exons could be detected. In these cases it is not known whether the truncation of the gene is a true biological event caused by recent recombination, or whether it is due to a mis-assembly of the genome sequence.

The optimised gene set was combined with the gene predictions in Ensembl (Methods, hand-filtered alignments and location checks of the remaining genes to identify accordance). The final list of novel zebrafish NLR proteins contains 368 members (Supplementary Excel Table1). A further 36 predictions for NLR genes had been annotated as pseudogenes and were therefore not retrieved for this list (online supplementary material). The refined genes have since been integrated into the VEGA and Ensembl gene sets, however since the annotation was performed on pre-GRCz10 paths, the latest GRCz10 gene set (Ensembl80) might differ marginally from the described results.

### Conserved NLR genes across metazoa

We used the zebrafish gene identifiers for the conserved NLRs in zebrafish to query the Orthoinspector 2.0 database (Linard et al. 2014) at http://lbgi.igbmc.fr/orthoinspector for orthologs in published genomes and downloaded the corresponding sequence. We then queried a custom Blast database of the Cyprinus carpio proteome, as well as the NCBI nr database for selected fish species using BLASTP. After removing redundant hits we calculated alignments employing Clustal-Omega v.1.2 (Sievers et al. 2011) and subsequently removed sequences of poor quality. In a second inference we also used trimal (Capella-Gutiérrez et al. 2009) to reduce the alignment to the conserved residues. We employed prottest v.3.2 (Darriba et al. 2011) to infer the best fitting evolutionary model and found that the LG model with Gamma optimisation performed best under the AKIKE information criterion. We then ran RAxML v7.7.2 (Stamatakis 2006) on both alignments on the Cologne University CHEOPS super computer and calculated bootstrap values. Phylogenetic trees were visualised and edited in Dendroscope v.3.2.5 (Huson et al. 2007).

### figmop and tblastn screen for NLR-B30.2 candidates in other fish genomes

Expanded gene families are not well annotated in most genomes. Rather than relying on gene predictions for identifying NLR-B30.2 genes, we therefore directly searched the genome sequences of 6 species: *Latimeria chalumnae, Lepisosteus oculatus, Callorhinchus milii, Esox lucius, Astyanax mexicanus, Cyprinus carpio*. We downloaded genome data either from NCBI servers or the genome project websites. We then used the Figmop (Curran et al. 2014) pipeline to find contigs and scaffolds in the genomes with NLR-B30.2 candidates on them. The Figmop pipeline builds a profile of conserved motifs from a starting set of sequences and uses these to search a target database with the MEME software suite (Bailey et al. 2009).

We used zebrafish NLR-B30.2 sequences from all four groups to create a set of 15 motifs to earch the above genomes. The resulting contigs were then subjected to the Augustus (v3.0.3) gene prediction pipeline (Stanke and Waack 2003) to predict genes *de novo*, setting zebrafish as the “species”. We complemented this approach by TBLASTN searches using the NACHT as well as the B30.2 domains as queries in individual searches and then kept those predictions in which the domains occurred in the proper order (thereby excluding spurious cases caused by mis-assembly or incomplete genes).

### Recursive phylogenetic analysis

We used a recursive approach for identifying genes for the phylogeny that were representative of the overall sequence divergence in the gene family. We selected only those that had both the NACHT and B30.2 domains. We then recursively performed the following: (1) constructed a sequence alignment of the sequences in the dataset using Clustal-Omega, (2) constructed a phylogeny using a Bayesian approach using MrBayes with mcmc=1,000,000, sump burnin=1000 and sumt burnin=1000 and (3) removed monophyletic paralogs from the dataset. The recursive analysis was halted when no instances of paralogous sister-sequences remained (Fig. 3), with the exception that at least one zebrafish and carp sequence from each of the major groups was retained. Once the final dataset of sequences was determined we removed gap-containing and highly variable columns from the alignment and re-ran MrBayes with mcmc=2,000,000, sump burnin=2000 and sumt burnin=2000 and re-confirmed our inferred tree with maximum likelihood in RAxML.

### Divergence analysis

For zebrafish genes in groups 1, 2a, 3a and 3b that contain both NACHT and B30.2 domains we calculated all pairwise ds values using PAML (Yang 2007). For each such gene we recorded the comparison with the smallest ds across all 4 groups. For genes in one group we then calculated average smallest ds values (rows in Table 2).

## ACKNOWLEDGEMENTS

We thank Robert Remy and Giuliano Crispatzu for some early work on this project and Jonathan Wood for verifying genomic locations of assembly components. Financial support was provided by EMBO and the DFG SFB 670 “Zellautonome Immunität” to ML, DFG SFB 680 “Molecular basis of evolutionary innovation” to TW, the HHMI International Early Career Scientist Program (55007424), MINECO (Sev-2012-0208), AGAUR program (2014 SGR 0974), and an ERC Starting Grant (335980_EinME) to FK, the European Molecular Biology Laboratory to JM, the Wellcome Trust to KH (zebrafish genome sequencing project) and the National Human Genome Research Institute (NHGRI) grant HG002659 to GKL (gene annotation), and a grant from the Volkswagen Foundation to PHS. We thank the CHEOPS support team and the Bundesland Nordrhein Westfalen for making HPC applications freely available at the University of Cologne.

## Supplementary Figure Legends

**SFig 1:**
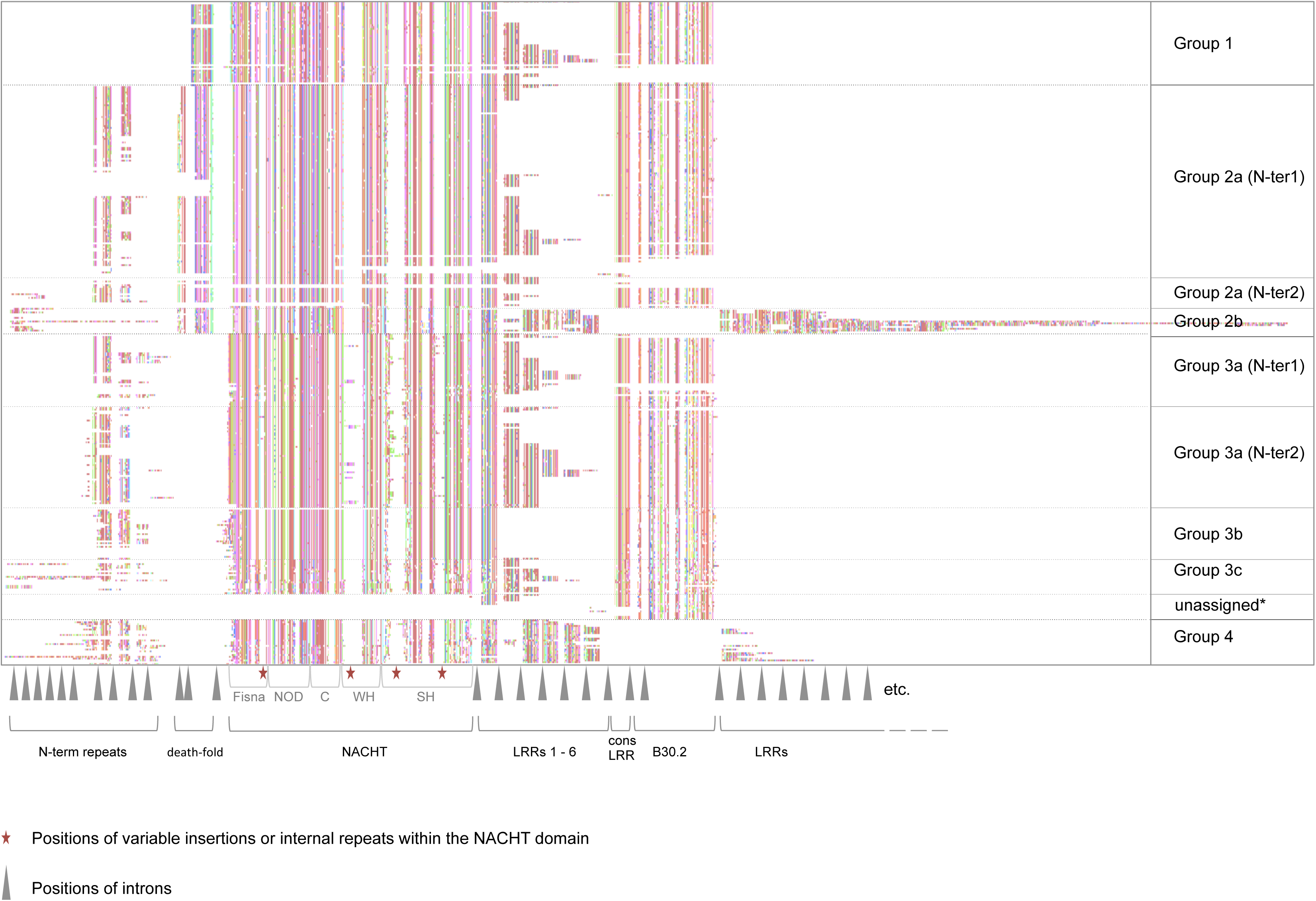
Overview of the entire set of 368 predicted NLR-B30.2 proteins in the zebrafish, based on a Clustal Omega alignment. The original alignment (online supplementary material) was edited by hand to improve the alignments of the N-terminal repeats and the LRR. The colour code for the amino acids was assigned at random. Gaps were introduced in the alignment at the positions of introns (marked by a grey arrow head below the alignment). Further large gaps are created because some positions are prone to variable and often long insertions or internal duplications (marked by red asterisks below the alignment). The domains are marked below the sequences; domain boundaries within the NACHT domain are entered according to reference (Proell et al. 2008). Positions of introns are marked by grey arrowheads below the sequences.

**SFig 2:**
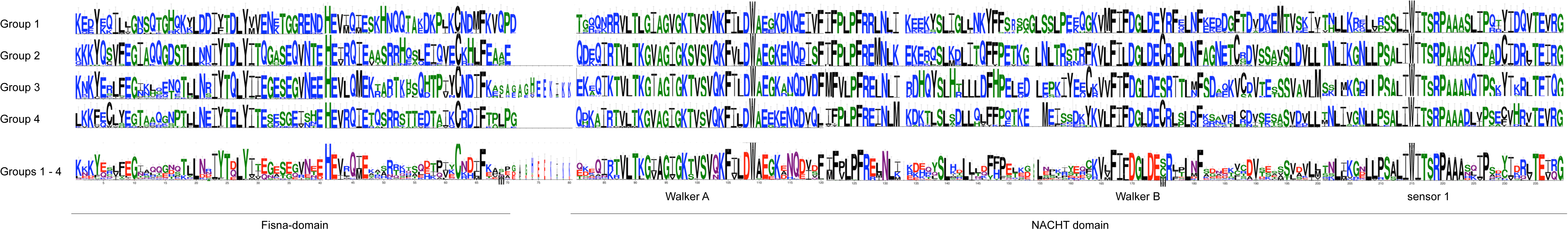
Comparison of FISNA and NACHT domains in groups 1 – 4 in the NLR-B30.2 genes. Logos generated in Jalview from the alignment in the online supplementary material. Differences in the Walker – A motive are diagnostic for groups.

**SFig 3:**
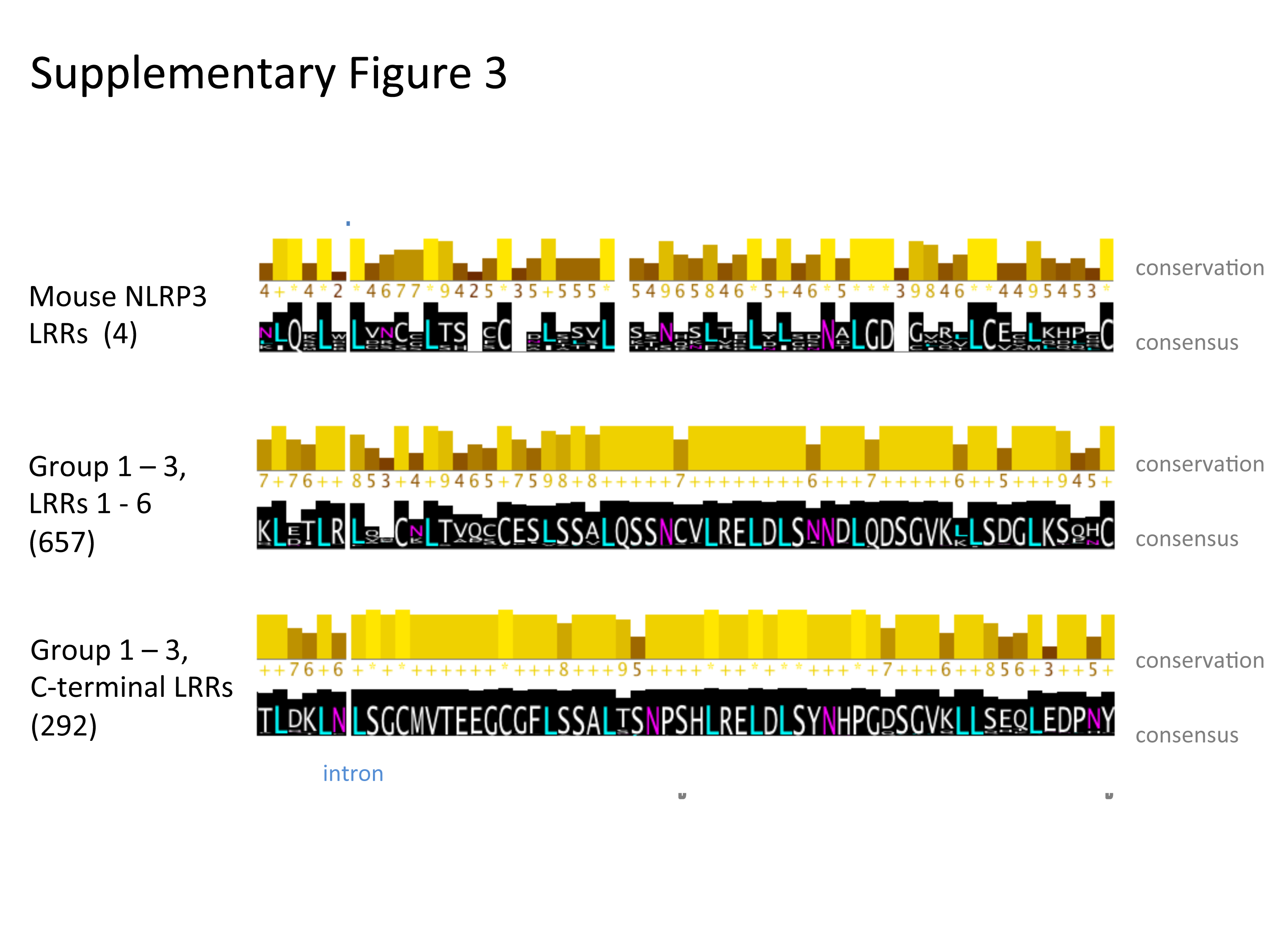
Comparison of LRRs in groups 1 – 3 and mouse NLRP3 LRRs. Logos generated in Jalview from the alignment in online supplementary material.

**SFig 4:**
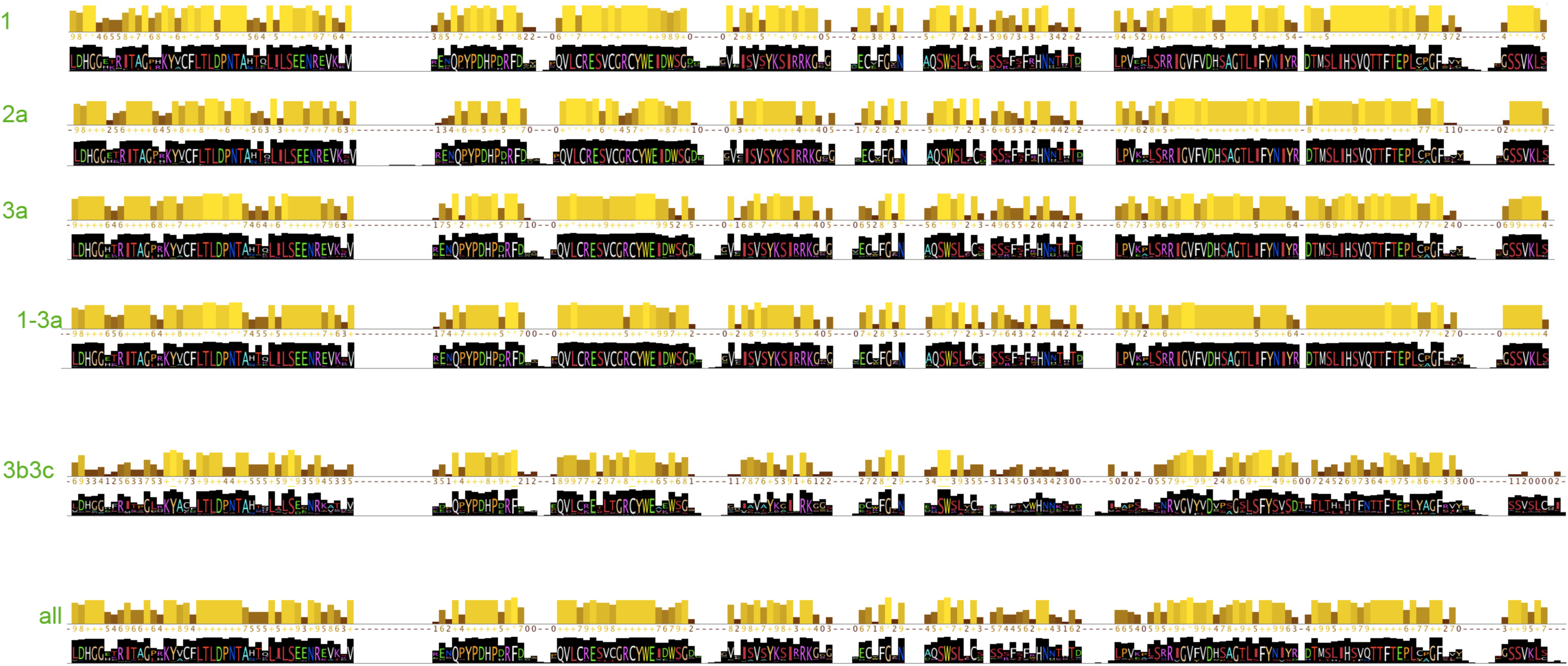
Differences in the B30.2 domain between the indicated group. Logos generated in Jalview from the alignment in online supplementary material.

**SFig 5:**
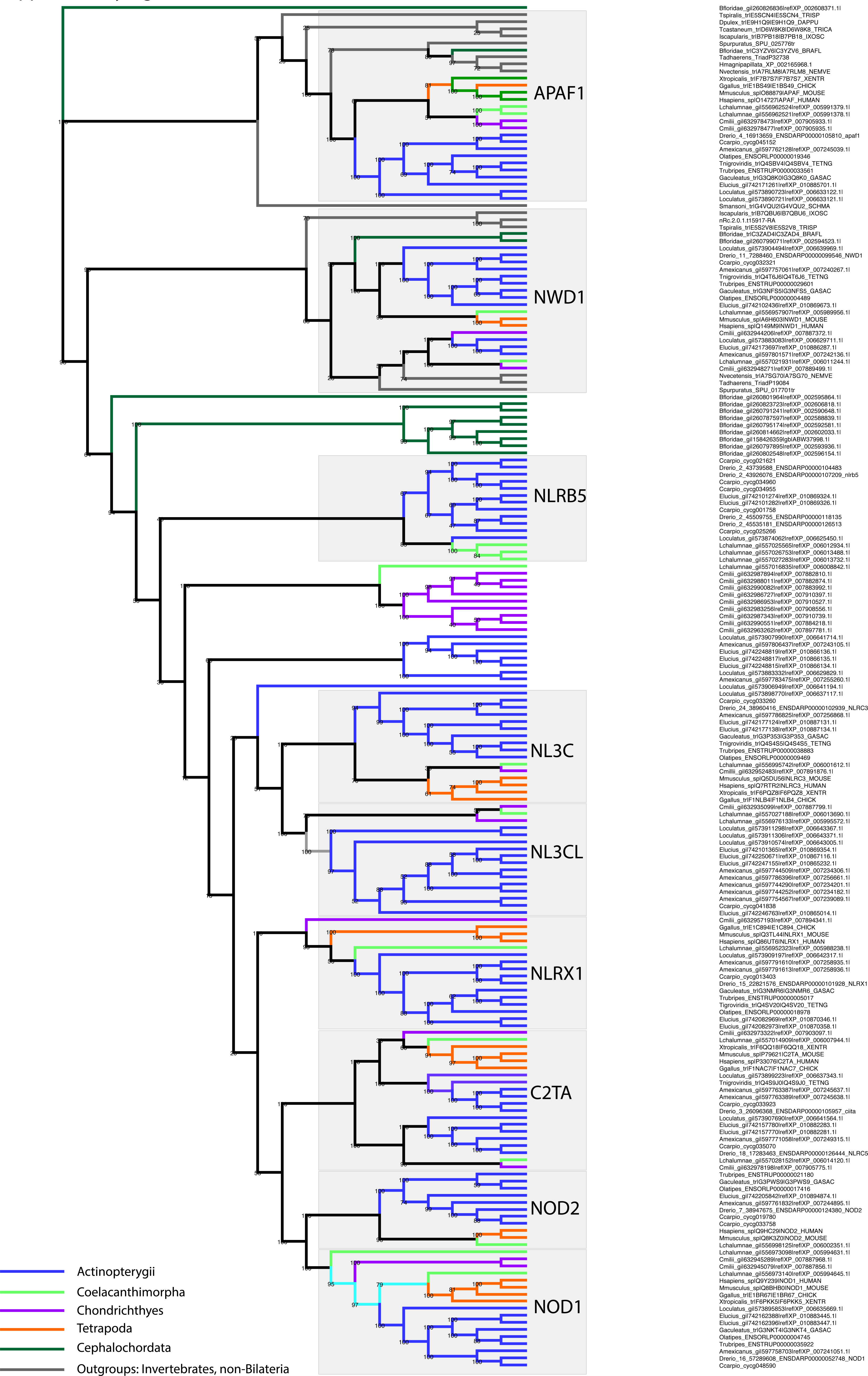
Phylogenetic ML tree of the conserved NLR proteins in Metazoa.

